# Source Reconstruction of Resting-State MEG and EEG Activity: A Technical Note on the Choice of Noise Covariance

**DOI:** 10.1101/2025.08.20.671381

**Authors:** Alexander Moiseev, Sam M. Doesburg, George Medvedev, Vasily A. Vakorin

## Abstract

Minimum variance beamforming is widely used to reconstruct neural sources from MEG and EEG data, but results critically depend on the choice of noise covariance. In task-based studies, this is often defined from pre-stimulus baselines, but for resting-state data the problem presents a fundamental challenge. Conventional solutions, such as empty-room recordings or diagonal white sensor noise, are not optimal. They either ignore brain-generated noise or yield artificial, non-uniform source-level baselines that can distort results. Our approach is to define a baseline at the source level as a uniform distribution of uncorrelated, randomly oriented neural dipoles, representing a maximum-entropy “ground state” of brain activity. Projecting this source model through the electromagnetic lead fields yields a sensor-level covariance that captures realistic spatial correlations. A data-driven constraint scales the model to match measured data, ensuring a physically admissible solution. Applied to real human resting-state data, the method produces a structured, non-uniform sensor covariance dictated by participant’s anatomy, source reconstructions that are smooth and plausible, and free from the artificial peaks induced by diagonal noise models. This source-level approach provides a principled and physiologically grounded baseline for beamforming and improves the reliability of resting-state analyses and interpretation.

## 1. Introduction

Neuroimaging techniques such as magnetoencephalography (MEG) and electroencephalography (EEG) provide an unparalleled non-invasive window into the temporal dynamics of human brain function [1,2]. By deploying an array of sensors around the head, these methods capture the weak electromagnetic fields generated by synchronous neural activity. However, the data recorded at the scalp or just outside it are a complex superposition of signals from multiple underlying neural sources. The fundamental challenge is to reconstruct the spatial distribution and temporal evolution of electromagnetic activations within the brain volume from these external measurements. This is known as the bioelectromagnetic inverse problem, a classic ill-posed problem for which no unique solution exists without the imposition of additional constraints and modeling assumptions [1,3].

Among the various techniques developed to provide an approximate solution, the minimum variance beamformer, which has been known for quite some time in radar field and acoustics [4], over the last two decades has emerged as a popular tool within the neuroimaging community [5–7]. A beamformer is a linear spatial filter designed to reconstruct the time course of activity from a specific target location in the brain while optimally suppressing signals originating from all other locations. This paper will focus on the scalar formulation of the minimum variance beamformer, which models individual brain sources as current dipoles with fixed positions and orientations. The filter weights are calculated individually for each source and directly depend on these parameters. However in nearly all practical applications source locations and orientations are not known; therefore, the application of beamforming necessitates a preceding source localization step.

The nature of this localization problem differs significantly between stimulus-driven and resting-state paradigms. In stimulus-based studies, the goal is to identify focal and spatially compact brain activations time-locked to a task. Localization is achieved by scanning the source parameter space with a “localizer function”, which typically is some form of a signal-to-noise ratio (SNR). Maxima of the unbiased localizer correspond to the true source parameters. In the simplest so-called “single source” approach all activations of interest are treated as independent [7,8]. In other scenarios several simultaneous and possibly correlated activations must be considered together and more complex multi-source localizers must be applied [1,9–11]. Importantly, in both those situations the noise covariance matrix is a critical component of the localizer construction. For stimulus based studies the noise covariance can be empirically estimated from pre-stimulus or “baseline” intervals where the signal of interest is absent.

In this paper our focus is on resting-state activity, which is the spontaneous, ongoing neural dynamics of the brain in the absence of any explicit task. In the resting state, it is assumed that continuously active sources are distributed across a predefined grid spanning the brain volume or the cortical surface. The localization problem thus shifts from finding the position of a few focal sources to determining optimal orientations for a vast number of distributed sources. While the same localizer functions can be applied, it becomes unclear which noise covariance matrix should be used for their construction. There is no obvious “control” or “baseline” intervals from which it can be estimated, although attempts to identify such intervals have been made [12].

The lack of a clear method to define the noise covariance is the central problem we aim to address. We should also take into account that since the measured sensor covariance matrix ***R*** is a sum of the source signal covariance and the noise covariance, ***N***, our choice of “noise” also dictates what will be reconstructed as a “signal”. Ideally the noise should be determined individually for each brain source, and should encompass signals from all other yet unknown brain sources, in addition to environmental and instrumental noise. In reality this is very hard to achieve, although such attempts have been made [13]. Using simulated or surrogate resting state data as a baseline was also suggested [14]. However in practice simplified, and often flawed, noise models are most commonly used.

The most popular approaches currently employed in resting state studies are twofold, and in our view neither is adequate. The first approach is to use an “empty room” covariance matrix, recorded with no subject present. While this method correctly captures the instrumental and environmental noise characteristics of the M/EEG system, these factors constitute only a minor part of the total interference at the source level. The dominant source of interference for any single brain location is the collective activity of all other brain sources. By using an empty-room covariance, this major contribution to the noise field is completely ignored.

The second common approach is to assume a diagonal white noise model, where ***N*** is proportional to an identity matrix (see for example [15–17]). Often this is a built-in default option in popular software toolboxes. Intuitively, diagonal white noise might appear to be a reasonable choice, representing a “uniform, uninformed prior” at the sensor level. However, this intuition is deeply misleading. A diagonal sensor covariance does not correspond to a uniform state at the source level. As we will show, it typically corresponds instead to a subject-specific, non-uniform, and spatially correlated distribution of neural sources. The exact pattern is determined by the subject’s head shape, the precise spatial positions of the sensors, and the parameters of the electromagnetic forward model. There is no physiological or theoretical reason why such a distribution should serve as a baseline condition for resting-state analyses.

In this technical note, we argue that in the absence of any *a priori* information, a more natural and principled baseline condition must be defined at the source level, not the sensor level. We propose that the most logical null hypothesis for resting-state brain activity is a uniform distribution of uncorrelated and randomly oriented brain sources. This represents a “ground state” from which organized, synchronous network activity can emerge as a statistically significant deviation. From this source-level model, the corresponding sensor-level noise covariance matrix can be derived by projecting the electromagnetic fields of these sources to the sensor array. As we demonstrate, the resulting matrix is in general neither diagonal nor resembles the covariance of white noise. We show that this noise covariance matrix can be constructed based on a simple expression and then properly scaled using a data-driven constraint to ensure the mathematical and physical consistency of the beamformer solution. Finally, we demonstrate the application of this approach to a real example of human resting-state MEG data.

## 2. Methods

We begin by formalizing the relationship between the underlying neural sources and the signals measured by the MEG/EEG sensor array, formulate the scalar minimum variance beamformer solution, and proceed with source localization and a critical role of the noise covariance. We then derive the expression for the proposed noise covariance matrix and explain how the “ground state” source power should be set based on the recorded data.

### 2.1. The M/EEG Measurement Model and Beamformer Solution

We assume that a recording is performed by an array of *m* sensors. The data at any given time point *t* can be represented by an *m*-dimensional column vector of sensor readings, ***b***(*t*). This measured field is modeled as a linear superposition of signals generated by *n* distinct neural sources and an additive noise term, ***ν***(*t*). Each source *i* is characterized by a set of parameters ***θ***^*i*^ and a time-varying amplitude *s*_*i*_(*t*). For the case of a current dipole model, these parameters consist of the source’s 3D location ***r***^*i*^ and its orientation, a unit vector ***o***^*i*^, such that ***θ***^*i*^ = {***r***^*i*^, ***o***^*i*^}.

By combining the time courses of all *n* sources into a single *n*-dimensional column vector ***s***(*t*) = {*s*_1_(*t*), *s*_2_(*t*), …, *s*_*n*_(*t*)}^*T*^, where superscript *T* denotes transposition, and corresponding forward solutions into an *m* × *n* lead field matrix ***H***(**Θ**) = {***h***(***θ***^1^), …, ***h***(***θ***^*n*^)}, we can express the standard bioelectromagnetic forward model as:

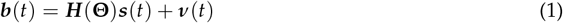

In Eq.(1) we assume that single source lead fields ***h***^*i*^ are known, and are static over the duration of the recording. Furthermore we assume that ⟨***ν***(*t*)⟩, ⟨***b***(*t*)⟩ = 0, where the angle brackets ⟨.⟩ denote statistical averaging, and that the processes ***s***(*t*), ***ν***(*t*) are stationary and uncorrelated with each other. The noise term ***ν***(*t*) conceptually encompasses all components of the measured field not produced by the *n* sources of interest, including instrumental noise and environmental interference.

The goal of the inverse problem is to reconstruct the source time courses ***s***(*t*) from the measurements ***b***(*t*) [1,3,7]. The beamformer approach achieves this by constructing a linear spatial filter for each source. Assuming the source parameters **Θ** are known, the linear filter estimate for the time courses of a subset of *k* ≤ *m* sources ***ŝ***(*t*) is given by applying a *m* × *k* weight matrix ***W*** to the data:

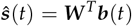

For the Multiple Constrained Minimum Variance (MCMV) beamformer, the weight matrix ***W*** is derived by minimizing the total output power of all *k* sources subject to the constraint ***W*** ^*T*^ ***H*** = ***I***_*k*_, where ***H*** = {***h***^*j*^} is the (*m* × *k*) lead field matrix for the selected sources and ***I***_*k*_ denotes *k*-dimensional identity matrix [1,4,9]. This yields the well-known multi-source beamformer solution:

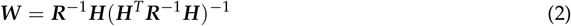

In Eq.(2) ***R*** = ⟨***bb***^*T*^ ⟩ is the (*m* × *m*) data covariance matrix, and columns ***w***^*j*^, *j* = 1, …, *k* of matrix ***W*** represent the weight vectors of individual sources.

In the multi-source formulation any change in the parameters of one of the sources affects the entire matrix ***W*** making all weight vectors ***w***^*j*^ interdependent. This interdependence significantly complicates the search for localizer extrema [9,10,18]. For this reason a simpler single-source approximation (*k* = 1) is most commonly used where each source is treated independently. In this case the expression for the weight vector ***w***^*i*^ of a single source *i* simplifies to:

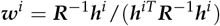

where now ***w***^*i*^ only depends on this source’s parameters ***θ***^*i*^.

The above solutions provide the time course estimates for a sources if their locations and orientations are known. The central issue addressed in this paper is the localization step that must precede this reconstruction, particularly the challenge of defining the noise model required for this step in the context of resting-state data.

### 2.2. The Localization Problem and the Critical Role of Noise Covariance

The expressions for the beamformer weights in Equation (2) are predicated on knowing the source parameters **Θ** – that is, the locations and orientations of the neural activity. In practice, these parameters are the very unknowns we wish to find. The process of determining them is the localization step. This is typically achieved by searching the parameter space for maxima of a so-called localizer function. For a localizer to be effective, its maxima should correspond to the true source parameters, providing an unbiased estimate of the source locations and orientations regardless of the signal-to-noise ratio (SNR).

In the context of resting-state analysis, where sources are assumed to exist across a predefined grid of positions ***r***^*i*^, the localization problem simplifies to finding the optimal orientation ***o***^*i*^ at each location. Historically many localizer functions have been proposed that were applicable only in a single-source case; and a few – for the multi-source case [5,6,8–11,18–20]. One of the most widely used and well understood is the pseudo-Z statistic [6,7], which for a single source comes down to the ratio of the estimated source power to the power of the noise projected by the spatial filter. This can be expressed as:

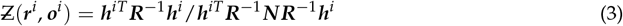

By construction, the pseudo-Z statistic is directly related to the source-level SNR (specifically, 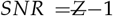). The optimal orientation ***o***(***r***^*i*^) at a given location ***r***^*i*^ is the one that maximizes this function. The resulting spatial map of 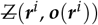 is then used to characterize the distribution of brain activity.

Equation (3) makes the fundamental challenge explicit: localization requires not only the measured data covariance ***R*** but also a separate, well-defined noise covariance matrix ***N***. This is a general situation, not restricted to this particular form of the localizer or to the single-source approximation. The choice of ***N*** is crucial, as it provides the baseline against which the signal is defined and detected. In stimulus-based paradigms, this is straightforward; ***N*** is empirically estimated from “control” or pre-stimulus time windows where the signal of interest is absent, and ***R*** is estimated from “active” or post-stimulus windows. In the resting state, however, no such control intervals exist. The entire recording is a continuous stream of the brain activity we wish to characterize, making the definition of ***N*** a non-trivial modeling decision that directly shapes the inverse solution.

### 2.3. A Principled Noise Model from a Source-Level Null Hypothesis

As argued in the *Introduction*, common choices for ***N*** in resting-state analysis, such as diagonal white noise or empty-room recordings, are physiologically and mathematically inconsistent. To overcome these limitations, we propose a more natural alternative by defining a baseline or null condition at the source level itself. We posit that a natural “ground state” for spontaneous brain activity is a uniform distribution of uncorrelated and randomly oriented dipolar sources distributed throughout the brain volume.

We formalize this model with the following assumptions for the baseline state:

1. **Uniform Source Distribution:** Sources *s*_*i*_(*t*) exist at all *n* locations ***r***^*i*^ within the source space.
2. **Uncorrelation and Equal Variance:** The sources are uncorrelated and share a common root mean square (RMS) amplitude, *σ*. This is expressed as ⟨*s*_*i*_*s*_*j*_⟩ = *σ*^2^*δ*_*ij*_, *i, j* = 1, …, *n*, where *δ*_*ij*_ is the Kronecker delta.
3. **Random Orientation:** The orientations ***o***^*i*^ are uniformly and independently distributed over the unit sphere. The expectation of the orientation outer product is therefore ⟨***o***^*i*^ ***o***^*iT*^ ⟩ = (1/3)***I***_3_, where ***I***_3_ is the 3 × 3 identity matrix.

The observed brain field ***b***(*t*) in this baseline or ground state is the superposition of the fields from all these sources, plus any additional environmental or sensor noise ***ν***(*t*):

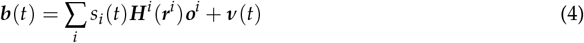

Here, we have introduced the *m* × 3 lead field matrices ***H***^*i*^ (***r***^*i*^) = {***h***^*ix*^, ***h***^*iy*^, ***h***^*iz*^}, which represent the sensor responses to unit dipoles at location ***r***^*i*^ oriented along the Cartesian axes. The noise ***ν***(*t*) is assumed to be a zero-mean process with covariance **Λ** = ⟨***ν***(*t*)***ν***(*t*)^*T*^⟩, which can be estimated from empty room recordings.

The noise covariance matrix ***N*** for our model is the expected covariance of this baseline field, ***N*** = ⟨***b***(*t*)***b***(*t*)^*T*^⟩. Using (4), straightforward algebra yields the following expression for our proposed baseline noise covariance matrix:

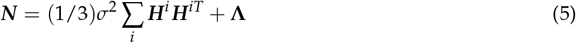

This model separates the contributions of physiology and anatomy from the overall source power. The term ∑_*i*_ ***H***^*i*^ ***H***^*iT*^ is a purely geometric construct that depends only on the head model and sensor layout. The scalar term (1/3)*σ*^2^ represents the uniform power of the underlying neural sources. The matrix **Λ** accounts for non-biological noise. This formulation provides a structured, non-diagonal, and physiologically motivated noise covariance that can be computed efficiently for any given source space. The final step is to determine the scalar source power *σ*^2^ in a data-driven manner, which we address in the following section.

### 2.4. Data-Driven Constraint and Model Scaling

The expression for the noise covariance matrix ***N*** derived in Equation (5) contains a single, crucial free parameter: *σ*, the root mean square (RMS) amplitude of the underlying, uniformly distributed neural sources. This parameter cannot be chosen arbitrarily, as its value has a direct physical interpretation and fundamentally impacts the resulting source reconstruction. To complete our model, we must establish a principled, data-driven method for determining its value based on the actual measured recordings.

To this end, we consider the relationship between the total measured sensor covariance, ***R***, and its constituent parts. ***R*** can be conceptualized as the sum of the covariance of our baseline “ground state” noise model, ***N***, and the covariance of the “activity of interest,” ***C***. The matrix ***C*** represents the organized neural dynamics – the deviations from the uniform, random baseline that we aim to detect and localize. This relationship is expressed as:

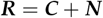

All three matrices – ***R, C***, and ***N*** – are covariance matrices, and are therefore symmetric and positive semi-definite. This reflects a critical physical principle that the power of any signal component cannot be negative. This fundamental requirement allows us to place an upper bound on the noise covariance and, consequently, on the value of *σ*. From ***C*** = ***R*** − ***N*** and constraint ***C*** ≥ 0 follows

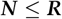

Here, an expression ***A*** ≤ ***B*** for two non-negative matrices ***A*** and ***B*** means that the difference ***B*** − ***A*** is also a non-negative matrix. Substituting our model for ***N*** from Equation (5), this constraint can be written explicitly as:

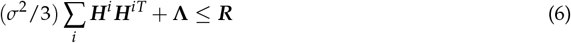

If the value of *σ* is set too high, this condition will be violated. A violation implies that the resulting signal covariance matrix ***C*** would no longer be positive semi-definite, which is a physically untenable situation that can lead to unphysical results, such as the estimation of negative source powers in certain brain regions. The constraint in Equation (6) therefore ensures the physical plausibility of the source reconstruction.

It is important to note that a value of *σ* that satisfies this constraint can always be found. When the measured covariance ***R*** is strictly positive, a sufficiently small *σ* will always satisfy the inequality. In practice, however, ***R*** may be rank-deficient due to common preprocessing steps such as the application of a common average reference in EEG, Maxwell filtering for noise cancellation in MEG, or the out-projection of specific interference components. This does not invalidate our approach. These same linear operations must be consistently applied to the components of our noise model, which will result in ***N*** having the same rank deficiency and null space as ***R***. Consequently, the constraint remains well-defined and can be evaluated within the common, non-null subspace of the data, ensuring the robustness of the method to standard preprocessing pipelines.

In practice, it is not only possible but also desirable to choose the maximal value for *σ* that still satisfies the constraint in (6). We denote this value as *σ*_*max*_. There are two primary reasons for this choice. First, when the environmental and instrumental noise **Λ** is not negligible, choosing *σ*_*max*_ ensures that the contribution of the brain’s own baseline activity to the noise covariance is maximized relative to the non-biological noise. This aligns with the fact that the dominant source of interference in MEG/EEG is typically the brain itself. Second, in the common case where **Λ** is negligible, choosing a value *σ* < *σ*_*max*_ merely leads to a proportional scaling of the pseudo-Z values without affecting the locations of the maxima. This increase in pseudo-Z is a purely artificial consequence of scaling down the denominator ***N*** and carries no additional physical significance. Selecting *σ*_*max*_ therefore provides a principled, data-driven approach to scaling the resulting activity maps.

The actual value of *σ*_*max*_ depends on the measured data ***R*** and the instrumental noise **Λ**, and it can always be found numerically using a straightforward iterative procedure to any desired accuracy. This completes the construction of our proposed noise model for beamformer analyses of spontaneous brain activity. The final, fully specified expression for the baseline noise covariance matrix is given by:

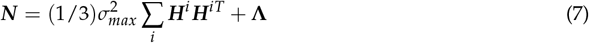

Importantly, equation (7) implies that ***N*** can never be diagonal because of the terms ***H***^*i*^ ***H***^*iT*^, even if the environmental noise covariance **Λ** is diagonal. Thus, white diagonal noise at the source level does not yield a white or diagonal covariance at the sensor level. Owing to the uniqueness of the electromagnetic forward solution, ***any source distribution producing a diagonal sensor covariance must be non-uniform and therefore cannot represent a flat baseline or a uniform prior*** – this is the central message of our technical note.

## 3. Practical example

### 3.1. MEG Data and Forward Model Construction

To demonstrate our proposed approach we used real human resting-state MEG data from a public repository [21,22]. The data were collected from a healthy 30-year-old male subject using a 275-channel CTF/VSM whole-head scanner housed in a magnetically shielded room. A total of five minutes of eyes-open resting state data were acquired. The raw data were preprocessed using third-order gradient noise cancellation, filtered to a 1–100 Hz band with 60 Hz power line notched out, and downsampled to a 200 Hz sampling rate. The subject’s power spectrum showed a prominent alpha-band peak (7–12 Hz) with a magnitude of about 8 dB.

The subject’s anatomical T1-weighted MRI was used to construct the forward model. A single-layer boundary element model (BEM) was generated to define the conductor volume, and an equally spaced volumetric grid with 5 mm resolution was created within this volume, resulting in approximately 13,700 source locations. Lead fields for each source were computed using the MNE Python software suite [23,24]. Given the effectiveness of the gradient compensation, the environmental and instrumental noise were considered negligible, and corresponding term (**Λ** in Equation (7)) was omitted for this demonstration.

### 3.2. Characterization of the Proposed Noise Covariance Model

The baseline noise covariance matrix ***N*** was constructed according to our proposed model, as defined in Equation (7). The full data covariance ***R*** was computed from the whole 5-minute recording, and the *σ*_*max*_ parameter was determined numerically to satisfy the constraint ***R*** − ***N*** ≥ 0. The observed effective sensor-level power signal-to-noise ratio, defined as *SNR* = *tr*(***R***)/*tr*(***N***) − 1, was approximately 8 for this dataset.

The resulting noise covariance matrix is visualized in Figure 1. As expected, it is immediately apparent that although the underlying source model assumes uniform and uncorrelated white noise at the brain level, the resulting sensor-level noise covariance ***N*** is far from diagonal. The colormap plot (Figure 1A) reveals strong non-diagonal elements, indicating widespread spatial correlations between sensors. This is a direct consequence of the physical reality of M/EEG recordings: the electromagnetic field from any single current dipole spreads across the sensor array, inducing correlated signals in multiple sensors. The eigenvalue spectrum of ***N*** (Figure 1B) further highlights this complexity; it is not flat, as would be expected for white noise, but spans several orders of magnitude, reflecting the rich spatial structure of the aggregate lead fields.

**Figure 1.**
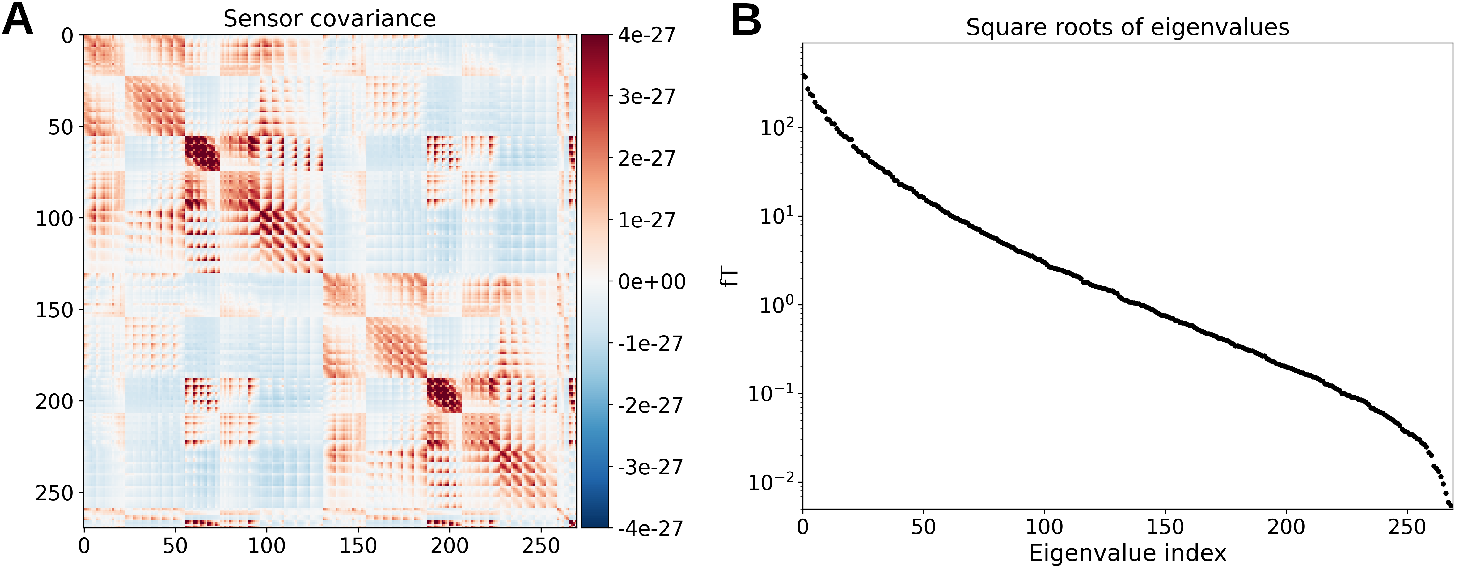
Noise covariance matrix constructed using model (7). *A* – Colormap of the matrix elements; channel indices are shown on the *X* and *Y* axes, and color values are in *T*^2^. *B* – Square roots of the eigenvalues of ***N*** (in fT). Note that only 270 out of 275 MEG channels were working.

Figure 2 depicts noise power distributions across the sensor array for both the recorded data and the model noise covariance. We notice that the original data exhibits strong activity in the parietal/occipital areas (Figure 2A), which is dominated by the alpha-band power (Figure 2B). Note also that the model noise power distribution is also highly non-uniform, as can be seen in Figure 2C. Its specific spatial pattern is not random; it is dictated by the subject’s unique head geometry, their position within the MEG helmet, and the parameters of the electromagnetic forward model. This again demonstrates that even a perfectly uniform source-level baseline produces a complex and highly structured sensor-level covariance. We also note a similarity between the distributions shown in Figure 2A and Figure 2C. This indicates that variations in signal power across the sensor array do not necessarily correspond to similar variations in the underlying source activity. In fact, this underlying activity may be relatively uniform.

**Figure 2.**
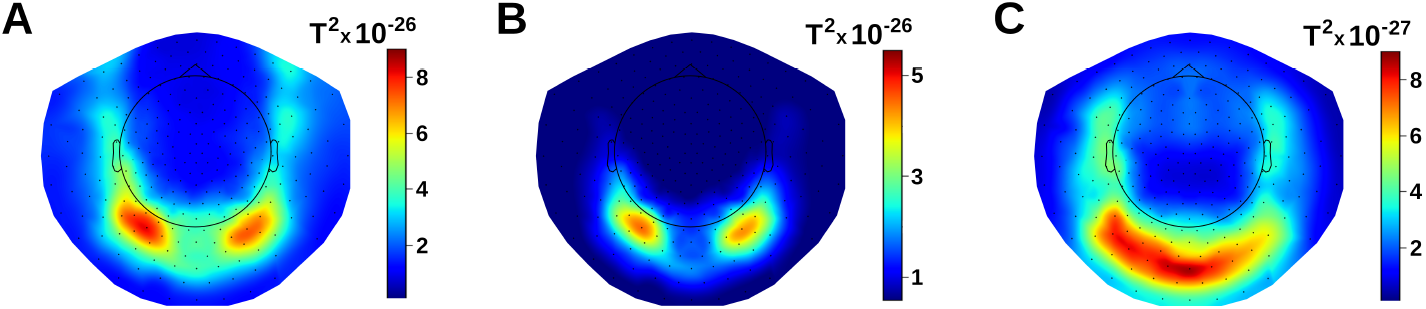
Topomaps of brain noise power distributions over the MEG sensor array. *A* – The real brain noise recorded in 1 to 100 Hz frequency range. *B* – The alpha band activity recorded in 7 to 12 Hz frequency range. *C* – The model noise power scaled to satisfy condition (6) for distribution in panel *A*.

### 3.3. Reconstructed Resting-State Activity: A Comparative Analysis

First of all, to demonstrate our main point regarding the conventional approach, we inferred how brain activity represented by the white diagonal covariance matrix would look like. Corresponding pseudo-Z distribution calculated using the true uniform baseline given by Equation (7) is presented in Figure 3A. This pseudo-Z map reveals a highly non-uniform and structured pattern, with pronounced peaks emerging in specific locations. This is the specific “activity” profile required at the source level to produce diagonal noise at the sensor level. Such a distribution is an unnatural and hardly justifiable choice for a baseline or “control” condition.

**Figure 3.**
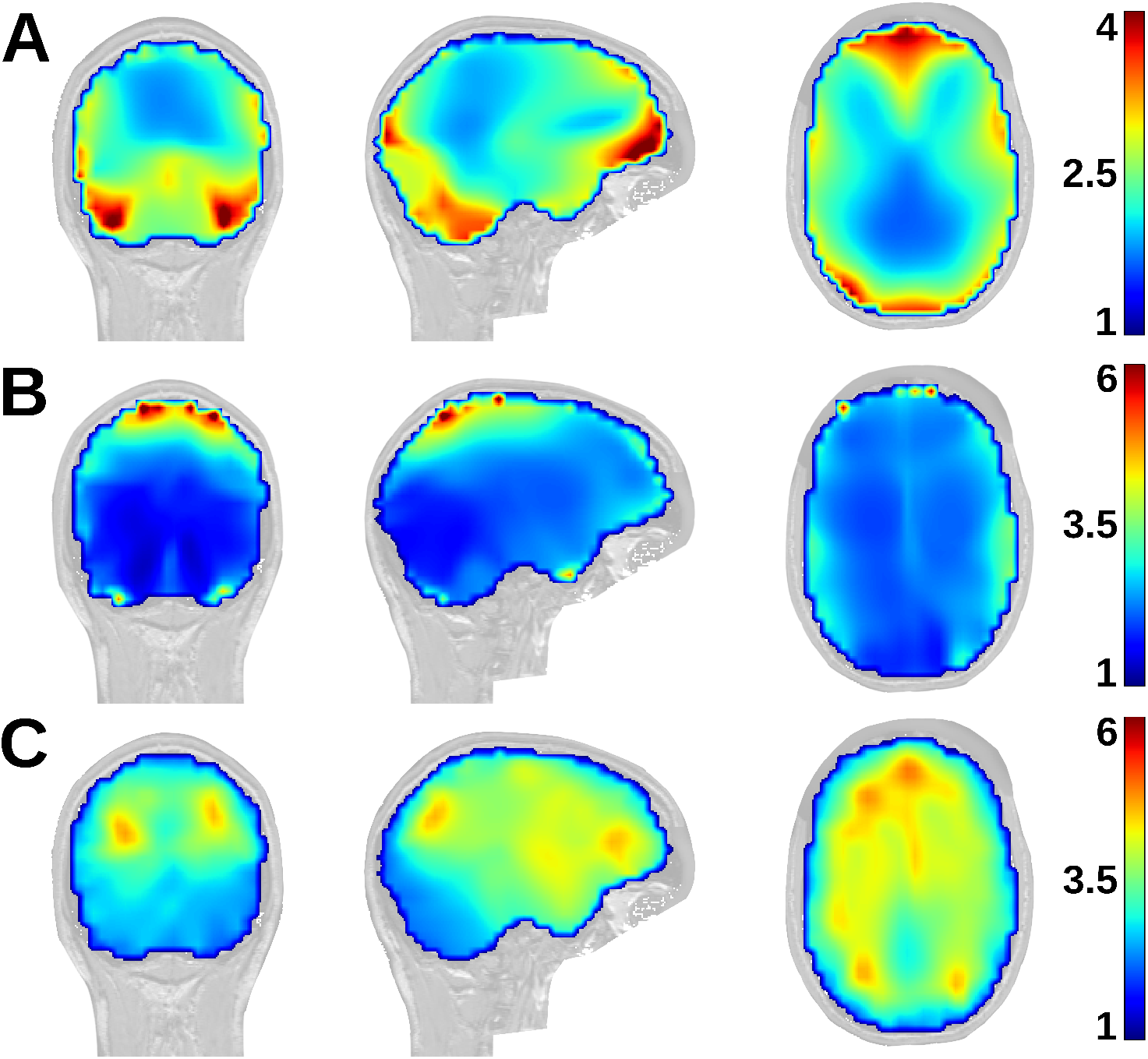
Pseudo-Z distributions of activity obtained for white sensor noise and for real brain noise in the 1–100 Hz range, computed using different baseline noise covariance matrices. Coronal and sagittal slices are taken approximately through the right-sided power peak in Figure 2A, while the axial slice is positioned midway between the vertex and the center of the head. *A* – diagonal white noise ***N***_*diag*_ versus the uniform baseline ***N*** defined by Eq. (7); *B* – real brain noise ***R*** versus the model baseline ***N***; *C* – real brain noise ***R*** versus the diagonal noise ***N***_*diag*_; ***N***_*diag*_ was scaled to satisfy constraint ***N***_*diag*_ ≤ ***R***.Note that the color bar limits for panel *A* differ from those for panels *B* and *C*.

We then applied our proposed model to compute the pseudo-Z distribution of brain activity corresponding to the observed real sensor covariance ***R*** (Figure 3B). This distribution exhibits a pattern consistent with neurophysiological expectations for resting-state activity. Pseudo-Z values – and thus SNRs – are higher for shallow sources and gradually decline with depth toward the center of the head. The distribution is spatially smooth, without sharp gradients or localized peaks, aside from the pronounced alpha clusters characteristic of this subject. Overall, the map provides a naturally looking representation of ongoing spontaneous resting-state brain activity.

Finally, we recomputed this pseudo-Z map of the actual measured data ***R*** using a scaled diagonal sensor noise matrix as the baseline, as is common in practice (Figure 3C). The result looks dramatically different from that obtained with our proposed model (Figure 3B). In particular, the map shows exaggerated frontal activity not present in Figure 2A, as well as artificially strong SNRs for deep sources near the center of the head. These effects arise solely from the non-uniformity of the chosen baseline and correspond to the spurious structure seen in Figure 3A. To further illustrate this, Figure 4 compares alpha-band pseudo-Z distributions obtained using our model noise versus diagonal white noise. As expected, alpha activity is strongest in parietal and temporal regions (Figure 4A). In contrast, the distribution based on diagonal sensor noise shows the largest pseudo-Z values in frontal and deep brain regions (Figure 4B). This comparison clearly illustrates how the conventional choice for the noise covariance can introduce systematic biases that could easily be misinterpreted if one mistakenly expects the white diagonal sensor noise to represent a flat baseline.

**Figure 4.**
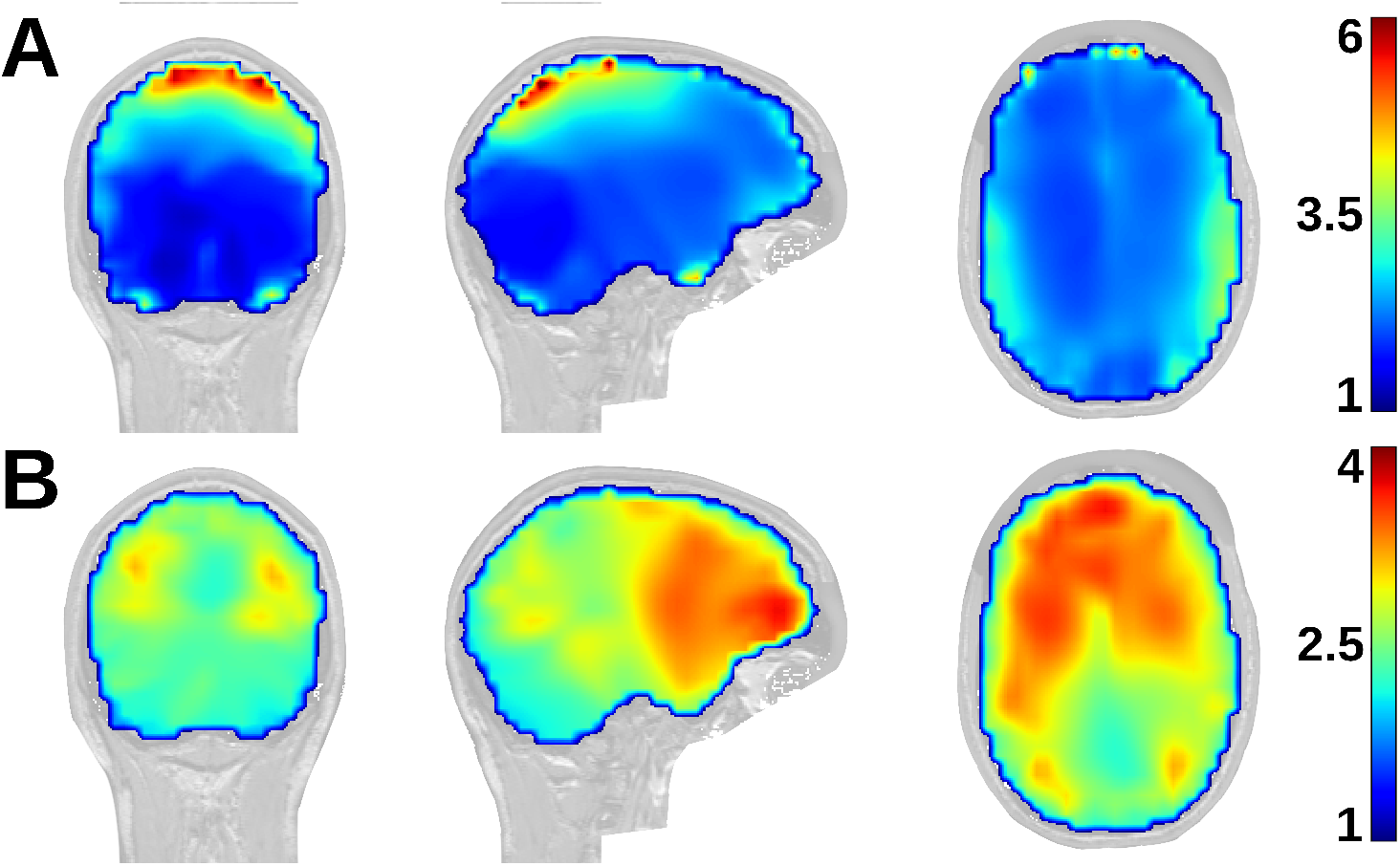
Pseudo-Z distributions of activity in alpha band (7 to 12 Hz), with the same slice coordinates as in Figure 3. *A* – the model brain noise ***N*** defined by Eq. (7) is used as the baseline; *B* – the diagonal noise ***N***_*diag*_ used as a baseline. Note that the color bar limits for panels *A* and *B* differ.

## 4. Discussion

The practical example in the preceding section illustrates that the choice of the baseline noise covariance matrix ***N*** is the very foundation upon which the entire beamformer analysis is built. The matrix ***N*** explicitly enters into the expressions for localizer functions [1,7], such as the pseudo-Z statistic in Equation (3), and is thus the primary determinant of the localization outcome. It is worth noting that while the final beamformer weights (Equation (2)) appear to depend only on the full data covariance matrix ***R***, they are indirectly shaped by the noise model. This is because the lead fields ***h*** depend on the source orientations that are themselves estimated during the localization step. Since localization is guided by the definition of noise, the choice of ***N*** ultimately affects both the estimated source parameters and the resulting spatial filters that determine the neural time courses.

In resting-state analysis, unlike in focal activation studies, the pseudo-Z distribution itself, rather than its extrema, is the quantity of interest. A pseudo-Z value reflects the expected SNR of the reconstructed signal, not the absolute strength of neural activity. Because both the covariance matrix ***R*** and the forward model ***H*** are fixed, the resulting pseudo-Z distribution is determined entirely by the choice of baseline noise *which is defined by the investigator*. Moreover, since the true baseline state of spontaneous activity cannot be established for an individual subject, *no “ground truth” exists for comparison*. This constitutes an inherent limitation of applying the beamformer approach to resting-state data analysis.

In such a situation, most researchers would prefer a flat reference when no other information is available. Diagonal sensor noise is often *assumed* to provide such a flat baseline, but Equation (7) shows that this assumption is incorrect (see also Figure 3A). It imposes a particular structure on the SNR estimation and does not correspond to a uniform reference. Moreover this baseline is physiologically untenable.

The consequence is that pseudo-Z maps constructed under this assumption can be misleading. Apparent differences between brain regions – such as stronger frontal than parietal alpha activity in our example in Figure 4B – may in fact arise from the choice of baseline rather than true underlying neural differences. Our main message is that this point is often overlooked, and that ***mistaking diagonal noise for a flat baseline can systematically bias the interpretation of resting-state beamformer results***. The same issue arises in EEG as well, and it remains regardless of whether a volumetric or cortical surface–based source space is used.

The proposed approach provides a plausible and simple-to-use null model against which true, organized network activity can be detected. However, we acknowledge that this model is a simplification and may not be optimal for capturing more complex or structured background activity that might be present in other contexts, such as during specific cognitive tasks or in certain clinical populations.

Finally, a practical note concerns the use of empty-room recordings to account for the instrumental and environmental noise component, **Λ**, in our model (Equation (7)). While often negligible, if this component is included, care must be taken when estimating *σ*_*max*_ using the constraint in Equation (6). In a real-world scenario, we work with statistical estimates of ***R*** and **Λ**, not their true, underlying values. Although in theory the difference ***R*** − **Λ** must be non-negative (since environmental and instrumental noise are components of the total measured signal), in practice this condition can be violated due to estimation errors. This is particularly likely if a significant amount of time has elapsed between the brain recording and the empty-room measurement, as the statistical properties of the environmental noise may have changed. In such cases, it is imperative to acquire updated measurements of **Λ** or to apply appropriate regularization techniques to ensure that the matrix ***R*** − **Λ** remains positive semidefinite before proceeding to determine *σ*_*max*_. This ensures the mathematical stability and physical plausibility of the final noise model.

## 5. Conclusion

In this technical note, we have addressed the problem of defining an appropriate baseline noise model for the application of minimum variance beamformers to the analysis of spontaneous brain activity. The beamformer is an inherently contrastive method; it interprets neural activity as a deviation from a predefined baseline state. Consequently, the proper definition of this baseline is not a mere technical detail but is essential for producing meaningful, interpretable, and valid scientific results. The choice of the noise model fundamentally shapes the inverse solution.

We argued that commonly used approaches for resting-state analysis are predicated on flawed assumptions. Methods such as employing empty-room recordings or assuming white, uncorrelated sensor noise lead to baseline models that, when mapped to the source space, correspond to highly non-uniform and subject-specific patterns of neural activity. Such baselines are difficult to justify, both methodologically and physiologically, as they introduce arbitrary and systematic biases into the source reconstruction. The resulting activity distributions are prone to misinterpretation.

As a more principled alternative, we proposed a model based on a physiologically plausible null hypothesis at the source level: a uniform distribution of uncorrelated, randomly oriented brain sources. This model serves as a “ground state” or maximum entropy state of brain activity. True signals of interest, such as the synchronous dynamics of resting-state networks, are then identified as statistically significant deviations from this uniform baseline.

Using real resting-state MEG data as an example, we demonstrated that the sensor-level covariance matrix associated with this source-level baseline model is profoundly different from the diagonal matrix of white noise. Our model naturally produces a highly structured covariance matrix that exhibits widespread inter-sensor correlations and a strongly non-uniform spatial power distribution, reflecting the underlying physics of MEG and the anatomy of the head. We further compared the reconstructed activity distributions obtained using our proposed model versus a conventional diagonal sensor noise model and found substantial differences in both the shape and magnitude of the results. While the ground truth in such data is inherently unknown, the activity map generated by our baseline model allows substantially more simple and physiologically plausible interpretation than artificially distorted map produced by the diagonal noise model.

Ultimately, this work provides a robust and theoretically grounded framework for defining the noise covariance in resting-state beamforming. By starting from a principled assumption at the source level, our approach avoids the arbitrary biases of conventional methods and yields a more easily interpretable view of the brain’s neural dynamics. This provides a more solid foundation for investigations into the complex functional architecture of the resting brain.

## Author Contributions

Conceptualization, methodology, software, original draft preparation, review and editing, A.M. and V.V.; review and editing, S.D. and G.M; supervision, project administration, funding acquisition, V.V. All authors have read and agreed to the published version of the manuscript.

## Funding

This research was funded by the Canadian Institutes of Health Research (CIHR), grant number 186270.

## Data Availability Statement

The data presented in this study were obtained from a public repository.

## Acknowledgments

This research was enabled in part by support provided by the Digital Research Alliance of Canada (*https://alliancecan.ca*).

## Conflicts of Interest

The authors declare no conflicts of interest.

## Notes

### Competing Interest Statement

The authors have declared no competing interest.

### Summary of Updates

The wording of the manuscript was improved to make the main message more clear. Real MEG data from a public repository is now being used as a practical illustrative example.

